# Elevated serum aspartate aminotransferase levels concomitant with normal alanine aminotransferase levels in older low body weight people: Preliminary findings from a community-based epidemiological study

**DOI:** 10.1101/528034

**Authors:** Michi Shibata, Kei Nakajima

**Affiliations:** School of Nutrition and Dietetics, Faculty of Health and Social Services, Kanagawa University of Human Services, 1-10-1 Heisei-cho, Yokosuka, Kanagawa 238-8522, Japan; Department of Endocrinology and Diabetes, Saitama Medical Center, Saitama Medical University, 1981 Kamoda, Kawagoe, Saitama 350-8550, Japan

**Keywords:** AST, ALT, BMI, low body weight, older people

## Abstract

**Background:** Serum enzyme levels, including hepatic transaminase, are unknown in older people with low body weight (LBW), who can easily experience sarcopenia. Therefore, we addressed preliminarily this issue in a cross-sectional study of an apparently healthy population.

**Methods:** We investigated the relationship of serum aspartate aminotransferase (AST), alanine aminotransferase (ALT), gamma glutamyl transpeptidase (GGT), alkaline phosphatase (ALP), lactate dehydrogenase (LDH), and total bilirubin levels with body mass index (BMI) and age in 79,623 subjects aged 20–80 years who underwent an annual checkup. The relationship between serum AST and serum creatinine, a surrogate marker of skeletal muscle mass, was also examined in 25,220 subjects who had data for serum creatinine.

**Results:** Serum levels of AST, ALP, and LDH levels were significantly higher in older (≥50 years) non-obese subjects compared with younger (< 50 years) corresponding subjects. Serum AST levels were significantly higher in older LBW subjects (BMI ≤18.9 kg/m^2^) than in those with a reference BMI of 20.9–22.9 kg/m^2^. Serum AST levels showed a J-shaped curve against BMI, whereas ALT and GGT levels showed a linear relationship, regardless of age. Serum levels of creatinine were significantly decreased across the increasing serum AST in men regardless of estimated glomerular filtration rate (eGFR) and women with eGFR ≥ 60 mL/min/1.73 m^2^ (p <0.0001).

**Conclusion:** Elevated serum AST levels concomitant with normal ALT levels, which might reflect systemic damage of skeletal muscle, may be prevalent in older LBW people. These conditions may have involved skeletal muscle damage. Further studies need to determine whether such a condition is equivalent to the etiology of sarcopenia.

## Introduction

Whether serum transaminase levels are decreased or increased in individuals with low-body weight (LBW) is unclear. In the geriatric population, LBW individuals are at increased risk for sarcopenia, an age-associated loss of muscle mass and function, which lead to disability, hospitalization, and death (Fielding et al. 2011, Calvani et al. 2015).

Aspartate aminotransferase (AST), alanine aminotransferase (ALT), and gamma glutamyl transpeptidase (GGT), which are routinely measured in an ordinary checkup, are present in multiple organs, including the liver, skeletal muscle, heart, and kidney (Lott and Landesman. 1984, Nathwani at al. 2005, Malakouti et al. 2017). However, ALT and GGT are predominantly found in the liver. Therefore, elevated serum AST levels concomitant with normal serum ALT levels could reflect injury of skeletal muscle and myocardium in clinical practice (Nathwani at al. 2005, Malakouti et al. 2017). Alkaline phosphatase (ALP) and lactate dehydrogenase (LDH) are also found in multiple organs, whereas ALP is preferentially included in bone tissue (Malakouti et al. 2017, Lowe and John. 2018). LDH is involved in similar organs, such the liver and skeletal muscle, where AST is found (Lott and Landesman. 1984, Nathwani at al. 2005, Malakouti et al. 2017).

We investigated the relationship of serum levels of AST, ALT, GGT, ALP, LDH, and total bilirubin, a marker of hepatic function (Gazzin et al. 2016), with body mass index (BMI) in an apparently healthy, general population aged from 20–80 years. To confirm that high serum AST with normal serum ALT can reflect skeletal muscle damage, the relationship between serum AST and serum creatinine, a surrogate marker of skeletal muscle mass (Hosten et al. 1990), was also examined.

## Methods

This preliminary cross-sectional study was part of an observational study that was performed to examine the relationships between lifestyle-related diseases and cardiometabolic risk factors (Muneyuki et al. 2013). The current study involved 2 institutions in Kanagawa and Saitama, Japan: Kanagawa University of Human Services and the Saitama Health Promotion Corporation, a public interest corporation. The protocol was approved by the Ethics Committees of Kanagawa University of Human Services (No.10-22). Informed consent was obtained from all patients who were included in the study.

### Subjects

Clinical data were obtained for 116,817 apparently healthy individuals, who underwent routine check-ups in Saitama Prefecture between April 2007 and March 2008. Inpatients and disabled individuals who could not move without assistance were not enrolled. After also excluding individuals with incomplete data of AST, ALT, and GGT, 79,623 subjects remained in the study (54,190 men and 25,433 women). To investigate the effect of age, subjects were divided into five age groups: 20–29, 30–39, 40–49, 50–59, and 60–80 years.

The relationship between serum AST and serum creatinine was examined in a subgroup of subjects who had data of serum creatinine (n = 25,220) after the restriction of subjects with serum ALT of < 30 U/L, which was conducted because of exclusion of high AST due to hepatic damage.

### Anthropometry and laboratory assays

Anthropometric measurements and the collection of blood samples for laboratory analysis were performed in the morning. BMI was calculated as body weight (kg) divided by height squared (m^2^). Subjects were divided into six BMI categories (≤18.9, 19.0–20.9, 21.0–22.9, 23.0–24.9, 25.0–26.9, and ≥ 27.0 kg/m^2^), as previously reported (Muneyuki et al. 2013). Because the prevalence of underweight subjects (BMI < 18.5 kg/m^2^) in their 50s and older was low (3.3%) in this study, we defined the lowest BMI category as ≤ 18.9 kg/m^2^, which was termed as LBW. When we selected these BMI categories, we took into consideration that the World Health Organization proposed that the BMI cutoff points for overweight and obesity in Asian populations should be ≥ 23.0 and ≥ 27.5 kg/m^2^, respectively. These cutoff points are lower than those in Western countries (WHO Expert Consultation. 2004).

To evaluate the relationship between serum AST and serum creatinine, serum AST level was classified into three ranges, < 20 U/L, 20–29 U/L, and ≥ 30 U/L. The estimated glomerular filtration rate (eGFR) was calculated using the following equation: eGFR (ml/min/1.73m^2^) =194 × serum Cr–1.094×age–0.287 (if female)×0.739, where Cr denotes serum creatinine concentration (mg/dL) (Matsuo et al. 2009). The eGFR was divided into two groups: ≥ 60 and <60 ml/min/1.73m^2^, a criteria for chronic kidney desease because serum creatinine is elevated in patients with renal failure.

### Statistical analysis

Data in Figure are expressed as means◻±◻standard erros. Significant differences in parameteres between BMI categories were evaluated using analysis of variance (ANOVA). Significant differences in serum creatinine between three AST categories were also evaluated using ANOVA and post-hoc Bonferroni test. Differences between subjects with a provisional reference BMI of 21.0–22.9 kg/m^2^ (Muneyuki et al. 2013, WHO Expert Consultation. 2004) and other BMI groups, and differences between the five age groups were evaluated using the *post-hoc* Bonferroni test and the Mann–Whitney test. Statistical analyses were performed using SAS-Enterprise Guide (SAS-EG 7.1; SAS Institute, Cary, NC, USA). P< 0.05 was considered to represent statistical significance.

## Results

All serum enzymes, except for ALP, in subjects aged ≥ 50 years and LDH in those aged ≥ 60 years, significantly increased across the increasing BMI categories (all P <0.0001, ANOVA; **Figure 1**). AST, GGT, ALP, and LDH levels were significantly higher in older (≥ 50 years) non-obese subjects (BMI < 25.0 kg/m^2^) compared with younger (< 50 years) corresponding subjects (Mann–Whitney test, all P < 0.0001). Notably, serum AST levels in older LBW subjects were significantly higher than those in subjects with a provisional reference BMI of 20.9–22.9 kg/m^2^ (P = 0.0003, Bonferroni test). Consequently, serum AST levels showed a J-shaped curve against BMI in older subjects. However, ALT and GGT levels showed almost a linear relationship against BMI, regardless of age groups. No significant difference was observed in serum total bilirbin levels between BMI and age groups.

**Figure 1.**
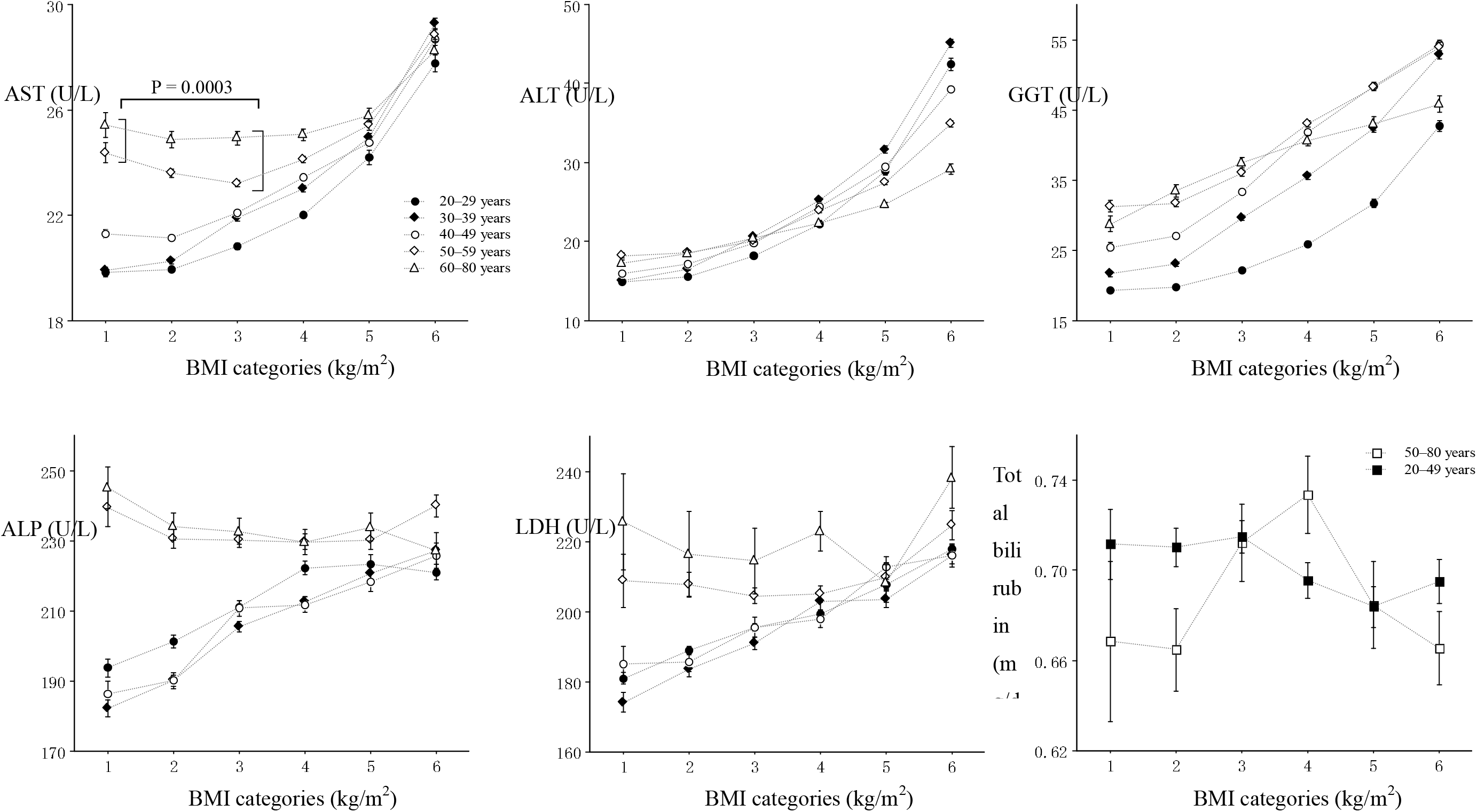
Serum enzyme and total birlibin levels according to BMI and age groups. The symbols indicate the mean values of parameters. The vertical bars represent the standard error. BMI categories of one to six represent ≤18.9, 19.0–20.9, 21.0–22.9, 23.0–24.9, 25.0–26.9, and ≥27.0 kg/m^2^. The number of subjects was 1848, 1745, 1268, 971, and 384 in 20s, 30s, 40s, 50s, and 60-80 years old, 3694, 3773, 3160, 2768, and 873 in 20s, 30s, 40s, 50s, and 60-80 years old, 4039, 4741, 4308, 4436, and 1541 in 20s, 30s, 40s, 50s, and 60-80 years old, 2683, 3712, 4229, 4627, and 1656 in 20s, 30s, 40s, 50s, and 60-80 years old, 1465, 2454, 2990, 3401, and 1141 in 20s, 30s, 40s, 50s, and 60-80 years old, 1564, 3065, 3465, 2935, and 687 in 20s, 30s, 40s, 50s, and 60-80 years old, for the BMI categories from one to six (AST, ALT, and GGT). Data of ALP, LDH, and total bilirubin were available only in 20,773, 7050, and 9061 subjects in total, respectively.

As shown in Figure 2, serum creatinine was decreased across the increasing AST in men regardless of eGFR and women with eGFR ≥ 60 mL/min/1.73 m^2^ (ANOVA, all p <0.0001). In Addition, Post-hoc Bonferroni test indicated that serum creatinine in subjects with AST ≥ 30 U/L was significantly lower than those with AST ≤ 19 U/L (all p <0.0001).

**Figure 2.**
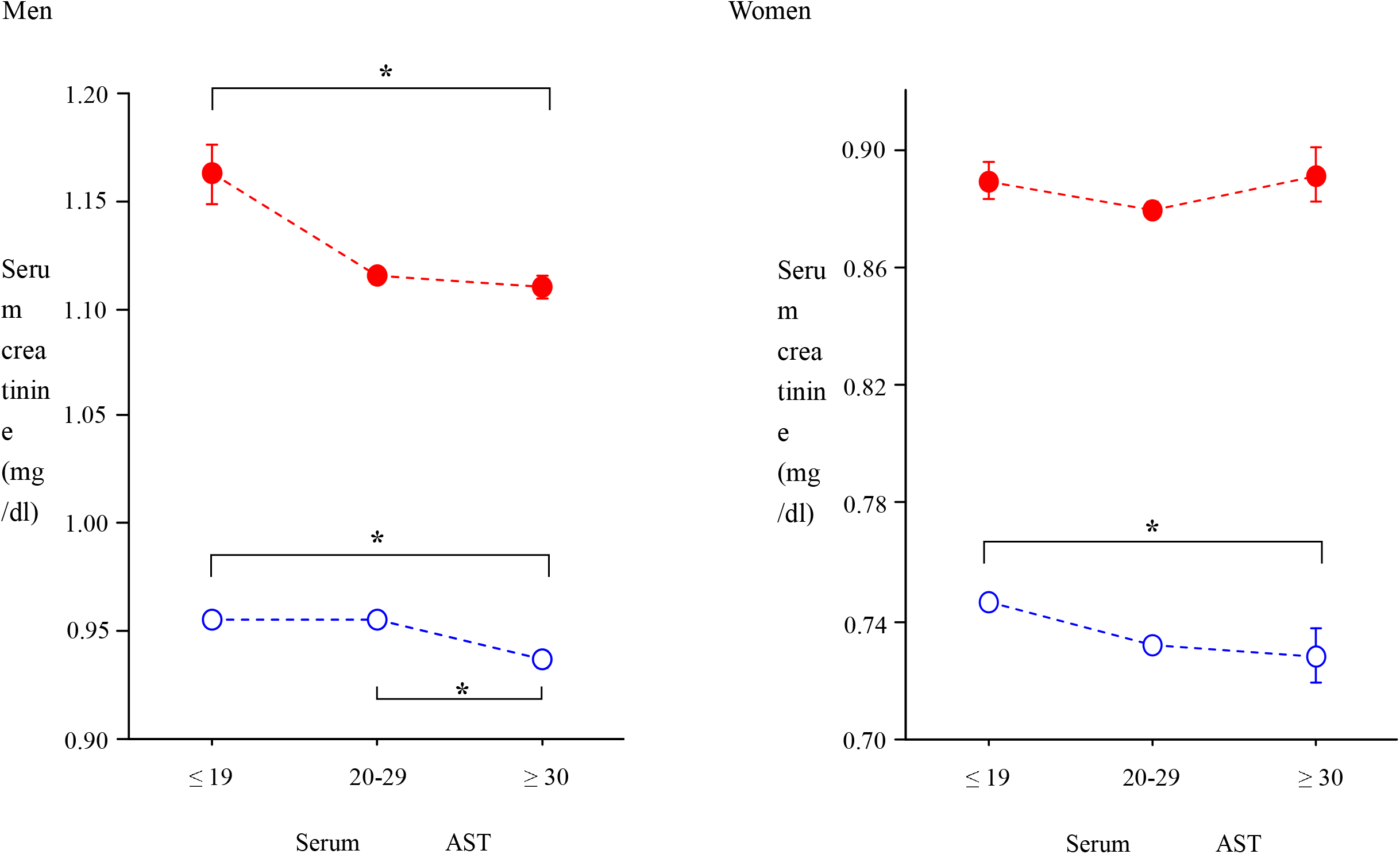
Serum creatinine levels according to three serum AST categories. The symbols indicate the mean values of parameters. The vertical bars represent the standard error. Open and closed circles represent subjects with eGFR ≥ 60 mL/min/1.73 m^2^ and subjects with eGFR < 60 mL/min/1.73 m^2^, respectively. * P <0.0001, Bonferroni test. The number of subjects was 3268, 4430, and 343 in men with eGFR ≥ 60 mL/min/1.73 m^2^, and 2500, 4629, and 411 in men with eGFR < 60 mL/min/1.73 m^2^, and 60-80 years old, and 2210, 1212, and 46 in women with eGFR ≥ 60 mL/min/1.73 m^2^, and 2987, 2967, and 217, for the three AST categories

## Discussion

To date, prevention of sarcopenia has been the focus of attention in aging societies (Fielding et al. 2011, Calvani et al. 2015). However, there is no blood marker for sarcopenia. A blood marker might enable us to detect sarcopenia earlier in the general population because the current definition of sarcopenia may be complicated and time-consuming to determine (Fielding et al. 2011, Calvani et al. 2015). The current preliminary study showed that LBW subjects in their ≥ 50s had elevated serum AST levels, which were acompanied by high ALP and LDH levels, but low to normal serum ALT levels, compared with coresponding younger subjects and those with a reference BMI. To the best of our knowledge, this is the first report to show such an observation. Although all enzymes measured in this study are likely present in multiple organs, ALT and GGT are predominantly found in the liver (Lott and Landesman. 1984, Nathwani at al. 2005, Malakouti et al. 2017). Therefore, elevated AST concomitant with normal ALT levels, resulting in a dissociation between serum AST and ALT levels in older LBW subjects, suggests non-hepatic injury. This could reflect systemic damage of skeletal muscle mass, including the myocadium. Some studies have reported that low ALT levels may be associated with sarcopenia (Ruhl and Everhart. 2013, Vespasiani-Gentilucci et al. 2018) This is partially consistent with our results because LBW subjects in our study had low to normal ALT levels compared with normal weight and obese subjects (Figure 1). High serum ALP levels in older LBW subjects may be attributable to concurrently occurring damage in bone tissue (Malakouti et al. 2017, Lowe and John. 2018). This possibility deserves further study. Unfortunately, isoforms of enzymes and serum creatinine kinase were unavailable in this study, which is a major limitation of this study. Additionally, other conditions that only elicit elevated AST levels were not thoroughly excluded.

Although circulating creatine kinase, aldolase, and myoglobin are indicators of skeletal muscle damage, these parameters are only measured in the clinical setting if a patient complains of muscle pain or the physician suspects myositis. Therefore, in the current study, we confirmed the inverse relationship between serum creatinine and serum AST in the subjects without high ALT, suggesting a rough trend that skeletal muscle mass is reduced in individuals with high AST and normal ALT.

In conclusion, older LBW people may have elevated AST comcomitant with normal ALT levels, compared with corresponding younger people and those with the reference BMI. These conditions may have involved skeletal muscle damage. Further study is required to determine whether this specific condition is equivalent to the etiology of sarcopenia.

## Acknowledgments

None

## Supportive foundations

None

## Author contributions

Shibata M and Nakajima K designed the study and analyzed the data. Nakajima K wrote the manuscript. Both authors have read and approved the final manuscript.

## References

Calvani R, Marini F, Cesari M, Tosato M, Anker SD, von Haehling S, Miller RR, Bernabei R, Landi F, Marzetti E; SPRINTT consortium. 2015. Biomarkers for physical frailty and sarcopenia: state of the science and future developments. J Cachexia Sarcopenia Muscle. 6:278–86. doi: 10.1002/jcsm.12051.

Fielding RA, Vellas B, Evans WJ, et al. 2011. Sarcopenia: an undiagnosed condition in older adults. Current consensus definition: prevalence, etiology, and consequences. International working group on sarcopenia. J Am Med Dir Assoc. 12:249–56. doi: 10.1016/j.jamda.2011.01.003.

Gazzin S, Vitek L, Watchko J, Shapiro SM, Tiribelli C. 2016. A Novel Perspective on the Biology of Bilirubin in Health and Disease. Trends Mol Med. 22:758–768. doi: 10.1016/j.molmed.2016.07.004.

Hosten AO. BUN and Creatinine. In: Walker HK, Hall WD, Hurst JW, editors. Clinical Methods: The History, Physical, and Laboratory Examinations. 3rd edition. Boston: Butterworths; 1990. Chapter 193.

Lott JA, Landesman PW. The enzymology of skeletal muscle disorders. 1984. Crit Rev Clin Lab Sci. 20:153–90. DOI: 10.3109/10408368409165773.

Lowe D, John S. Alkaline Phosphatase. 2018. StatPearls [Internet]. Treasure Island (FL): StatPearls Publishing; 2018-.

Malakouti M, Kataria A, Ali SK, Schenker S. 2017. Elevated Liver Enzymes in Asymptomatic Patients - What Should I Do? J Clin Transl Hepatol. 5:394–403. doi: 10.14218/JCTH.2017.00027.

Muneyuki T, Suwa K, Oshida H, et al. 2013. Design of the Saitama Cardiometabolic Disease and Organ Impairment Study (SCDOIS): A multidisciplinary observational epidemiological study. Open J Endocr Metab Dis. 2:144–56. DOI: 10.4236/ojemd.2013.32022

Matsuo S, Imai E, Horio M, et al. Revised equations for estimated GFR from serum creatinine in Japan. Am J Kidney Dis 2009;53:982–92.

Nathwani RA, Pais S, Reynolds TB, Kaplowitz N. 2005. Serum alanine aminotransferase in skeletal muscle diseases. Hepatology. 41:380–2. DOI: 10.1002/hep.20548

Ruhl CE, Everhart JE. 2013. The association of low serum alanine aminotransferase activity with mortality in the US population. Am J Epidemiol. 178:1702–11. doi: 10.1093/aje/kwt209.

Vespasiani-Gentilucci U, De Vincentis A, Ferrucci L, Bandinelli S, Antonelli Incalzi R, Picardi A. 2018. Low Alanine Aminotransferase Levels in the Elderly Population: Frailty, Disability, Sarcopenia, and Reduced Survival. J Gerontol A Biol Sci Med Sci. 73:925–930. doi: 10.1093/gerona/glx126.

WHO Expert Consultation. 2004. Appropriate body-mass index for Asian populations and its implications for policy and intervention strategies. Lancet. 363:157–63. DOI: 10.1016/S0140-6736(03)15268-3

